# The high diversity of *Scedosporium* and *Lomentospora* species and their prevalence in human-disturbed areas in Taiwan

**DOI:** 10.1101/2022.06.10.495564

**Authors:** Yin-Tse Huang, Yun-Chen Fan, Tsu-Chun Hung, Chi-Yu Chen, Pei-Lun Sun

**Author notes:** Corresponding author Correspondence: Pei-Lun Sun, MD, PhD, Department of Dermatology, Chang Gung Memorial Hospital, Linkou, Taoyuan, Taiwan.

## Abstract

*Scedosporium* and *Lomentospora* are important opportunistic pathogens causing localized or disseminated infection in humans. Understanding their environmental distribution is critical for public hygiene and clinical management. We carried out the first environmental survey in urbanized and natural regions in Taiwan. Overall, *Scedosporium* and *Lomentospora* species were recovered in 130 out of 271 soil samples (47.9%) across Taiwan. We morphologically and molecularly identified five *Scedosporium* species and one *Lomentospora* species. All four major clinical relevant species were isolated with high frequency, i.e. *S. apiospermum* (42.5%), *S. boydii* (27.5%), *L. prolificans* (15.0%), and *S. aurantiacum* (8.8%); two clinically minor species, *S. dehoogii* (5.6%) and *S. haikouense* (0.6%), had moderate incidence. These fungal species have high incidence in urban (48%) and hospital (67.4%) soil samples, and have limited distribution in samples from natural regions (5%). Multivariate analysis of the fungal composition revealed strong evidence of the preferential distribution of these fungi in urban and hospital regions compared to natural sites. In addition, strong evidence suggested that the distribution and abundance of these fungal species are highly heterogeneous in the environment; samples in vicinity often yield varied fungal communities. Our results indicated that these fungal species are prevalent in soil in Taiwan and their occurrences are associated with human activities. Hygiene sensitive places such as hospitals should be particularly aware of the high frequency of the clinical relevant species.

## 1. Introduction

*Scedosporium* and *Lomentospora* are two fungal genera that accommodate pathogenic species to immunocompromised and occasionally immunocompetent hosts. These fungal pathogens cause a broad range of clinical manifestation including colonization of respiratory tract, cutaneous infections, allergic reactions, and severe localized or disseminated mycoses. Mortality rates of infected hosts are generally high. In a meta-analysis of 264 infection causes, mortality rate was reported at an average of 22.2% in immunocompromised and of 14% in immunocompetent patients (1). In disseminated infections, mortality rate can be extreme as high as up to 80% (2). Treatments of *Scedosporium* and *Lomentospora* are challenging due to their low susceptibility to many current antifungal drugs such as 5-flucytosine, amphotericin B, and triazole drugs (2). In conjunction with their prevalence in the environment, *Scedosporium* and *Lomentospora* are imperative, emerging pathogens that call for research input in improving the diagnostic methods, developing effective therapies, and systematically surveying their distribution in the environment.

*Scedosporium* and *Lomentospora* are ubiquitous soil dwellers in temperate regions and are less frequently encountered in tropics (3, 4). Their distribution in the environment is often associated with nutrient-rich substrates such as agricultural and garden soils (3, 5), crude oil-contaminated soil (6), and manure of livestock, poultry or cattle (7–9). In addition, they can be isolated from low aerated and high osmotic substrates such as sewage sludge (10, 11), polluted pond bottoms (12), brackish water (13), coastal tidelands (14), and wastewater treatments (15, 16). Their prevalence in soil environments manifests in the cases of their occurrence in tissues of small wildlifes such as lizards (17), gerbils (18), donkey (19), and pigeons (20). In contrast, surveys of their occurrence in indoor water and air samples indicated that they are limited distributed in these samples (10, 12, 21, 22), while again, are abundant in the soil of potted plants (21). Due to their prevalence in human active regions, *Scedosporium* was proposed as an indicator species for anthropogenic impacts (23). Given the pathogenic nature of these fungi, their occurrence in the environment are of concern, especially in hospital and indoor settings, where they are posing potential hazard to people with chronic respiratory disease and/or immunocompromised (24, 25).

Different *Scedosporium* and *Lomentospora* species pose different clinical relevances. Four species, *S. boydii, S. apiospermum, S. aurantiacum* and *L. prolificans* are the most clinically relevant species that account for most infections in reported cases (26, 27). With a few case reports, *S. dehoogii* can cause subcutaneous and osteoarticular infections (28, 29). To effectively distinguish clinical species from other clinically minor species, recent efforts have made substantial taxonomic revisions and species delimitation of *Scedosporium* and *Lomentospora* based on their morphological and molecular distinction (3, 30, 31). To date, there are 13 distinct *Scedosporium* species: *S. americanum, S. apiospermum, S. aurantiacum, S. boydii, S. cereisporum, S. dehoogii, S. desertorum, S. haikouense, S. hainanense, S. minutisporum, S. multisporum, S. rarisporum, S. sanyaense*. Three species, *S. ellipsoideum*, and *S. fusoideum, S. angustum*, are hypothesized cryptic species in the *S. boydii* species complex but without provision of proper nomenclature requirements, and therefore invalid (32). *Lomentospora* accommodates two species including *L. prolificans* and a recently reported *L. valparaisensis* (33).

Research on their environmental distributions, compared to the clinical studies (e.g. pathology, diagnosis, antifungal resistance), have been less emphasized. To date, there were a few systematic surveys conducted in Australia (34), Europe (35, 36), South America (37, 38), Africa (39), and Southeast Asia (40, 41). These studies have revealed a high variation of species assemblies among samples, locations, and countries [see reviews such as Luplertlop (2018) (42) and Rougeron et al (2018)]. This, coupled with a few non-targeted environmental disclosures of these species in close proximity to human living habitats (5, 22, 43, 44), has emphasized the importance of recognizing these potential infection sources in the environment.

Studies of *Scedosporium* and *Lomentospora* species in Taiwan are limited. There were only a few clinical case reports (45–52) and an environmental survey (25). This study, therefore, aimed to survey the occurrence of these species in the soil environment across Taiwan. To assess the risk factor of these species in varied facilities, we compared the species distribution between natural and urbanized areas. We explicitly tested: (1) Do human-disturbed regions accommodate more fungal species than in natural areas? (2) Do hygiene-sensitive areas (e.g. hospital) harbor more fungal species compared to other urbanized regions?

## 2. Materials and Methods

### 2.1 Sampling sites

During 2014 to 2016, we collected soil samples from 79 locations across Taiwan (Table 1). Sampling sites were selected to represent different levels of human activities (urban and natural); in particular, we stressed the hospital samples as another sub-category in urban samples as we wanted to highlight the distribution of these potential opportunistic fungi in the hygiene sensitive places. In each site,1 – 14 sampling units were randomly chosen. For each sampling unit, five points with 10 – 20 cm intervals from the center were randomly selected to represent the fungal community of each unit. Soil was sampled with a metal spatula, sterilized with 70% ethanol between samples to avoid cross contamination. Soil was sampled at a depth of 7 – 10 cm beneath the soil surface, put in a sterile plastic bag and gently squeezed to mix; the pooled soil sample was then filled in a 50 ml sterile Falcon tube. Soil samples were stored at 4 °C until the fungal isolation.

**Table 1.**
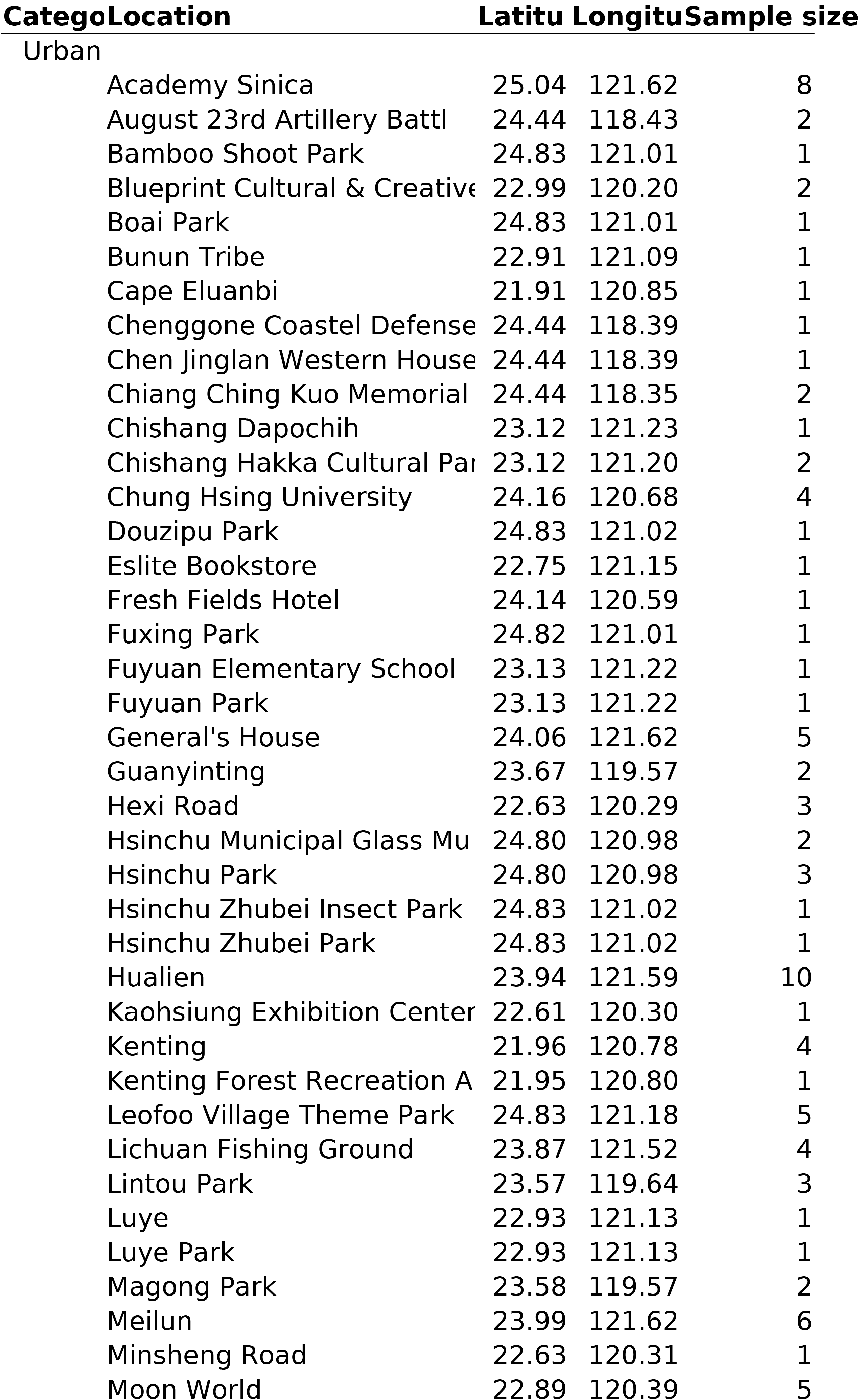
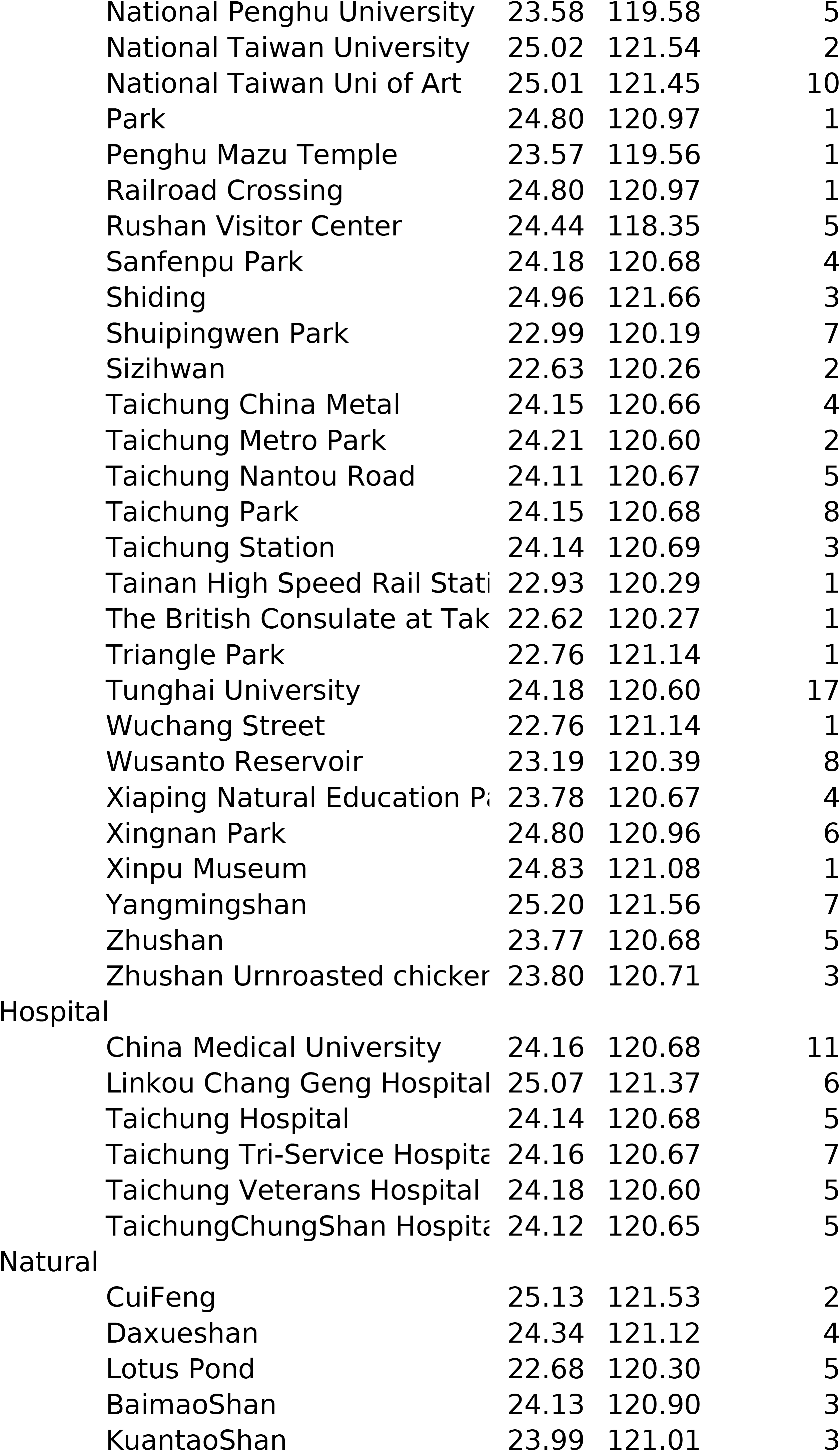
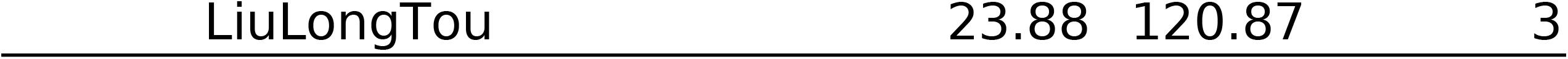
Sampling locations, assigned categories (urban, hospital, and hospital) of each sampling location, geographic coordinate, and sample sizes of each location.

### 2.2 Fungal isolation and fungal burden determination

For each soil sampling unit, ten grams of soil was air dried at room temperature for 1 – 2 days, placed in a 100 ml flask and homogenized with 90 ml sterilized water with 1 – 2 drops of Tween 20. The mixture was thoroughly vortexed and left to stand for 15 minutes. 250 μl of suspension of mixture were inoculated onto five plates of modified *Scedosporium* selective medium (SceSel+) (53), comprised of malt extract, 6.25 g; maltose, 6.25 g; mono-potassium-phosphate, 1.25 g; yeast extract, 1.0 g; magnesium sulfate heptahydrate, 0.625 g; soy peptone, 0.625 g; chloramphenicol, 0.1 g; streptomycin sulfate, 0.1 g; dichloran, 2 mg; benomyl, 6 mg; agar, 20.0 g; and reverse osmosis water, 990 ml. Plates were incubated at 37°C for 5 – 7 days and inspected at intervals for the emergence of characteristic colonies of *Scedosporium*. All *Scedosporium* colonies on five plates of each sampling unit were counted, from which the fungal burdens were determined based on the mean value of CFU per gram of soil dry weight.

### 2.3 DNA extraction, PCR, sequencing, and phylogeny reconstruction

Fungal tissues (5 – 20 mg) were lysed using mechanical beads (mix of 1 mm and 0.5 mm beads at 1:1 ratio) beating at 1800 rpm for 4 min using a Retsch mixer mill MM400. DNA extraction of the lysed cells were carried out using ZymoBIOMICS™ (CAT#D4306) with a ZiXpress 32 automated nucleic acid purification instrument following the manufacturer’s instructions. Quantity and quality of eluted DNA were measured using Qubit dsDNA BR Assay kit and Qubit Flex Fluorometer 4.0 (Thermo Fisher Scientific) following the manufacturer’s instructions.

For molecular identification, we respectively amplified the internal transcribed spacer (ITS) (samples started with “SUN”) and the full length of rDNA regions [including 18S small subunit (SSU), ITS, 28S large subunit (LSU)] for the rest of samples. Primer pairs and their references used for PCR amplification are shown in Table S1 (54, 55). Amplification was performed with a two-step barcoding approach according to Herbold et al. (56). The first-step PCR reaction mixture volume was 25 μl, consisted of 1 μl of DNA template, 1 μl of each primer, 12.5 μl of 2X Tools Supergreen PCR master mix (CAT#TTC-PA31-10), and 9.5 μl of ddH_2_O. Amplification were carried out as following steps: initial denaturation at 95 °C for 2 min, 25 cycles with denaturation at 95 °C for 30 s, annealing at 64 – 66 °C for 30 s, primer extension at 72 °C for 1 min, and final extension step was 7 min at 72 °C. The second-step PCR reaction mixture was 25 μl, consisted of 1 μl of the first-step DNA product as template, 2.5 μL pre-mixed barcoded primers, 10.75 μL of 2X Tools Supergreen PCR master mix, and 10.75 μl of ddH_2_O. The barcoded amplification products were purified using Ampure Xp Beads with 0.4X product to bead volume ratio, both procedures followed the manufacturer’s protocol. Sequencing was carried out on a Oxford Nanopore Technologies GridION sequencer using the manufacturer’s library preparation kit (SQK-LSK109).

We reconstructed the phylogeny of the isolated species using the ITS gene. We retried sequences of type specimens of *Scedosporium* and *Lomentospora* species from NCBI. Sequences of *Petriellopsis africana* (CBS311.72), *Petriella setifera* (CBS 385.87), and *Petriella sordida* (CBS 144612) were selected as the outgroup. We carried out the phylogenetic analysis using an in-house compiled workflow; sequences were aligned using MAFFT with automatic and adjustdirection flavor (57). Phylogenetic informative regions were selected using ClipKIT with gappy strategy (58). Phylogenetic trees were reconstructed using MrBayes (59) and RAxML-ng (60) with recommended partition parameters inferred using ModelTest-NG (61). The best model was GTR+I+G4 (Rmat= 2.1644, 3.3944, 2.5306, 1.1525, 6.2170, 1.0000; proportion of invariable sites=0.6736; gamma shape parameter=0.8810). For the MrBayes analysis, two MCMC runs of four chains were executed simultaneously from a random starting tree for 1000000 generations, every 500 generations were sampled resulting in 2000 trees, and 500 trees were discarded during burn-in. Posterior probabilities were estimated from the retained 1501 trees. Trees were visualized using Interactive Tree of Live version 4 (62).

### 2.4 Community analysis

We hypothesized that *Scedosporium* and *Lemontospora* species are more abundant in human-ditrubed regions than in natural regions. To test this hypothesis, we conducted a distance-based permutational multivariate analysis of variance (PERMANOVA) between the species distance matrix of urban and natural samples. This test measures if groups of observations differ in their composition. From our hypothesis, we expected to see a significant difference of species composition between urban and natural samples. The test was implemented on a Euclidean distance matrix using adonis() function of the vegan package for R (63). We then carried out a PERMANOVA pairwise post hoc test using pairwise.perm.manova() on the species distance matrix to test the differences of groups pairwisely (64). We further conducted a homogeneity test on the species community matrix using betadisper() function. Because the betadisper method is less sensitive to the variances among groups, it can help justify the significant result from PERMANOVA that is a result of compositional difference between samples, rather than within group variations. We presented Rsq and P-values and discussed their inferences using the language of “evidence” following the recommendation in Muff et al. (65)

To test for the effects of detecting *Scedosporium* and *Lomentospora* species in each location, location types (i.e. urban, hospital, and natural), and each sampling unit, we conducted a linear model-based analysis of the recovered species assemblage. We used the manyglm() function of the mvabund package of R (66) to fit individual negative binomial linear models for the species abundance of each detected fungal species. This approach provides an overall multivariate test of the effect of location and location type on the composition of the recovered fungal assemblage, as well as individual tests of the location, location types, and sampling units on the abundance of each fungal species with adjusted P-value to account for multiple comparisons. Three models were tested: (1) the effect of location type, (2) the effects of location type, location, and their interaction, and (3) the effects of location, sampling unit, and their interaction.

## 3. Results

### 3.1 Fungal isolation and fungal burden determination

160 *Scedosporium* and *Lomentospora* isolates were recovered from 271 sampling units in 79 locations, including 67 urban sites, six hospitals, and six natural sites across Taiwan (Table 1). The presence and abundance of these fungal species seems to be highly associated with human activities. They were detected in 48.0% of urban samples, 67.4% of hospital samples, and 5.0% of natural samples (Figure 1A). Likewise, the highest fungal burden was revealed in hospital samples with average 20.8 CFU/g, followed by urban samples with 10.29 CFU/g; in contrast, natural samples had only 0.11 CFU/g.

**Figure 1.**
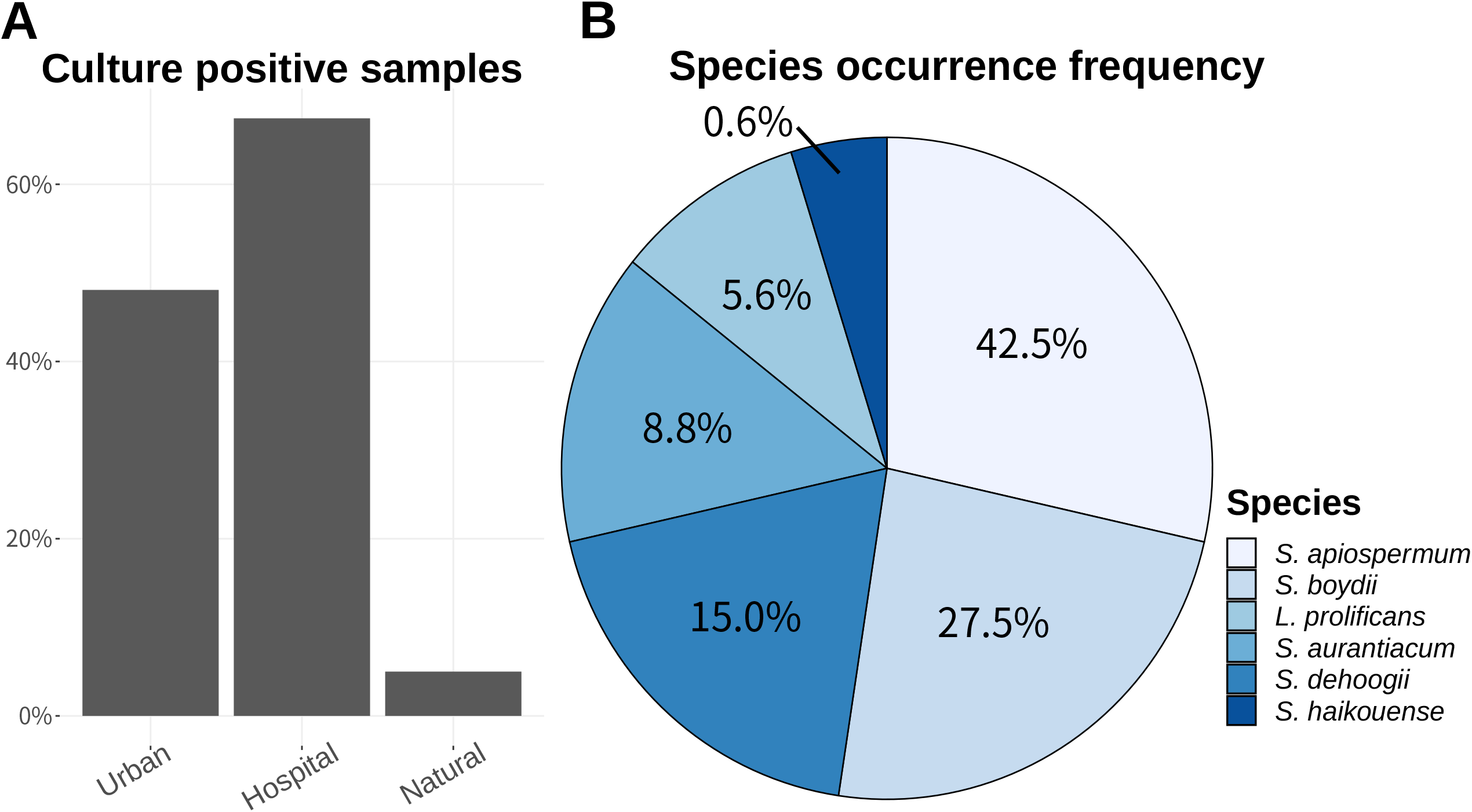
Culture-dependent isolation from soil samples in the present study. (A) Percentage of culture-positive samples of *Scedosporium* and *Lomemtospora* species in different types of sampling sources (urban, hospital, and natural) across Taiwan. (B) Proportion of each isolated *Scedosporium*/*Lomemtospora* species in the 164 isolates.

We identified six *Scedosporium* and *Lomentospora* species based on their morphological and molecular characteristics, they were *S. apiospermum, S. boydii, S. aurantiacum, S. dehoogii, S. haikouense*, and *L. prolificans* (Figure S1). Of all 160 isolates, the *S. apiospermum* was the most frequently recovered species (n = 68, 42.5%), followed by *S. boydii* (n = 44, 27.5%), *S. dehoogii* (n = 24, 15.0%), *S. aurantiacum* (n = 14, 8.7%), *L. prolificans* (n = 9, 5.6%), and *S. haikouense* (n=1, 0.6%) (Figure 1B). The occurrence of these fungal species among locations were highly varied (Figure 2 & 3). All six species were detected in urban samples, ranging from one species in 14 locations, two species in 11 locations, three species in 10 locations, four species in three locations, and all six species in Tunghai University. All hospital samples revealed the occurrence of these fungal species, ranging from one species in Linkou Chang Gung Memorial Hospital, two species in Taichung Tri-service Hospital, three species in four hospitals (Taichung Veterans Hospital, Taichung Hospital, Taichung Chung Shan Hospital, and China Medical University). In contrast, *S. apiospermum* was the only species recovered from the natural samples from Lotus Pond in Nantou.

**Figure 2.**
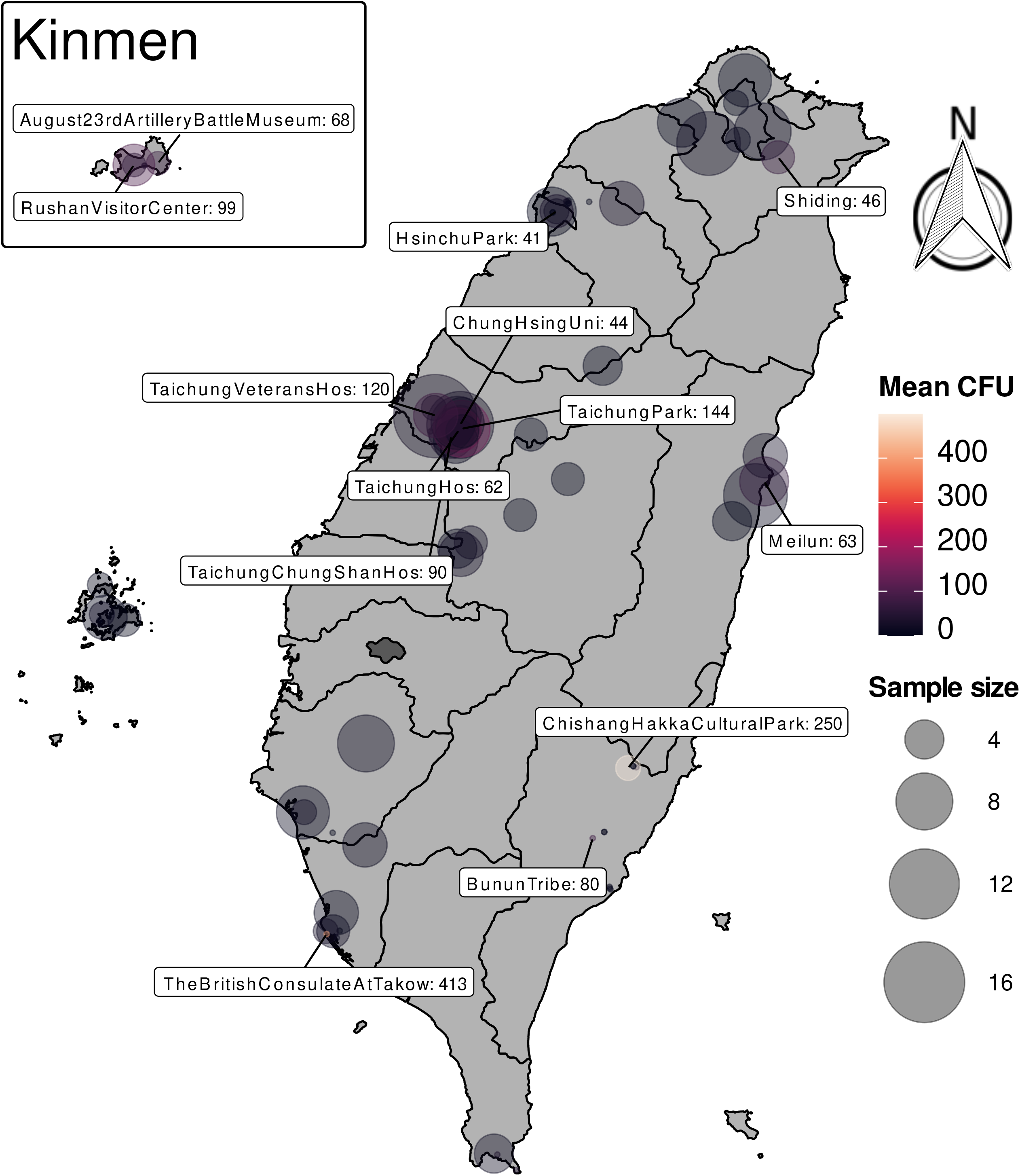
Mean CFU of *Scedosporium*/*Lomemtospora* species from samples in each location across Taiwan. Names of locations with mean CFU more than 40 are labeled. Circle sizes represent the sampling size in each location. * measured values of CFU/g.

Notably, the detected six fungal species exhibited some exceptionally high fungal burdens in certain sites. For example, high fungal burden of *S. apiospermum* were detected in six sampling locations including Chishang Hakka Cultural Park (499.5 CFU/g), Taichung Park (406.3 CFU/g), the British Consulate at Takow (365 CFU/g), Rushan Visitor Center (185.7 CFU/g), Taichung Hospital (185.1 CFU/g), and Meilun (112.0 CFU/g). *Lomentospora prolificans* was high in Taichung Veterans Hospital (406.7 CFU/g); *S. aurantiacum* was abundant in Rushan Visitor Center (182.8 CFU/g); and *S. dehoogii* showed its abundance the British Consulate at Takow (460 CFU/g) (Figure 3). The exceptionally high incidence of these opportunistic fungi on sites, especially for the clinically relevant species, raises the awareness to susceptible people such as immunocompromised and cystic fibrosis (CF) patients.

**Figure 3.**
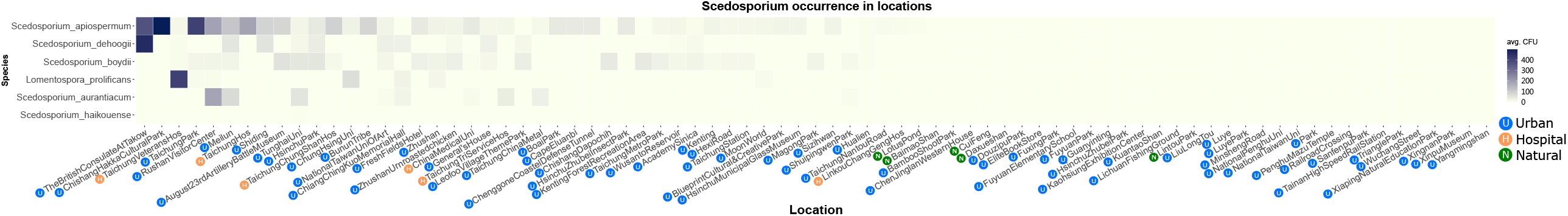
Heatmap showing the mean CFU of each *Scedosporium*/*Lomemtospora* species in each location sampled in the present study; types of sampling sources (urban, hospital, and natural) were labeled next the location name on x-axis.

### 3.2 Fungal community distribution analysis

Our analysis indicated there was an evident variation in the species composition of *Scedosporium* and *Lomentospora* within and between locations. Overall, there was strong evidence for the species composition in different location types (i.e. urban, hospital, and natural samples), while it accounted for a limited portion of the explained variation (adonis: df=2, *F*=5.299, *R*^*2*^=0.0351, *P*=0.002). While explaining much of the compositional difference, there was no evidence showing the species composition varied among locations (adonis: df=75, *F*=1.3057, *R*^*2*^=0.3247, *P*=0.033)(Table 2). In a pairwise PERMANOVA analysis, species composition in natural samples was distinct from that in urban and hospital samples, while no difference was detected between the urban and hospital samples (Table 3). The homogeneity test further indicated that species assemblage between natural-urban and natural-hospital samples was significantly different in their homogeneity; instead, no dispersion effect was found between urban and hospital samples (Figure 4) (betadisper: df=2, *F*=10.24222, *P*=5.1720e-05). The dissimilarity of species composition was also indicated by the clustering of natural samples, which were separated from most urban and hospital samples, on the NMDS plot (Figure 5). Altogether, these results suggested that *Scedosporium* and *Lomentospora* had different composition profiles in natural regions compared to human-dominated regions, i.e. urban and hospital samples.

**Table 2.**
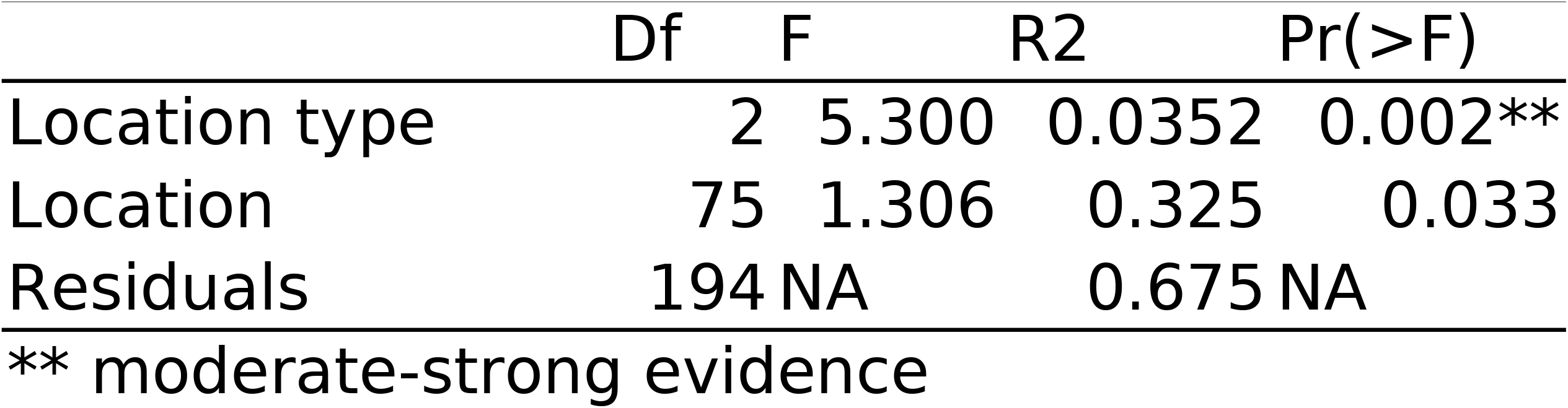
Degree of freedom, F-values, determination coefficients, and P-values of tested factors (location type and location) that influence the species assemblage observed in each location.

**Table 3.**
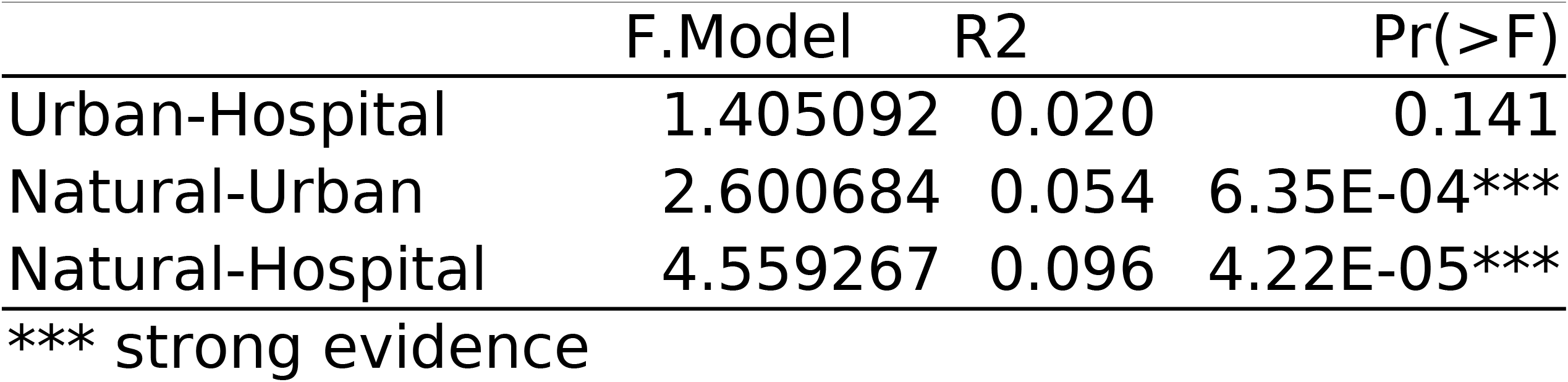
F-values, determination coefficients, and P-values of pairwise test of species assemblage observed in urban, hospital, and natural regions.

**Figure 4.**
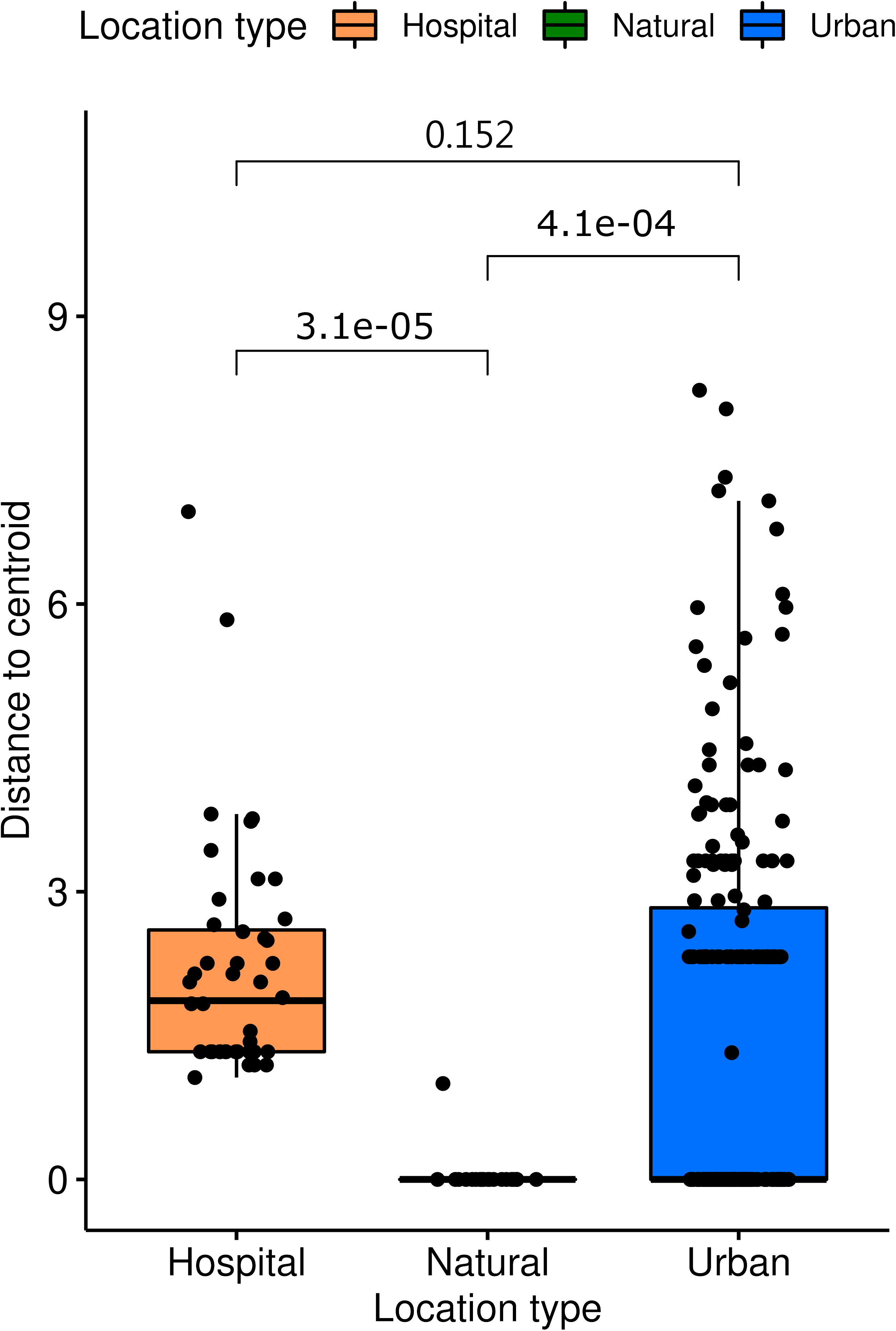
Homogeneity test on the species assemblage in different types of sampling sources (urban, hospital, and natural); the p-value generated via pairwise comparisons using post hoc t-test were shown above the bars.

**Figure 5.**
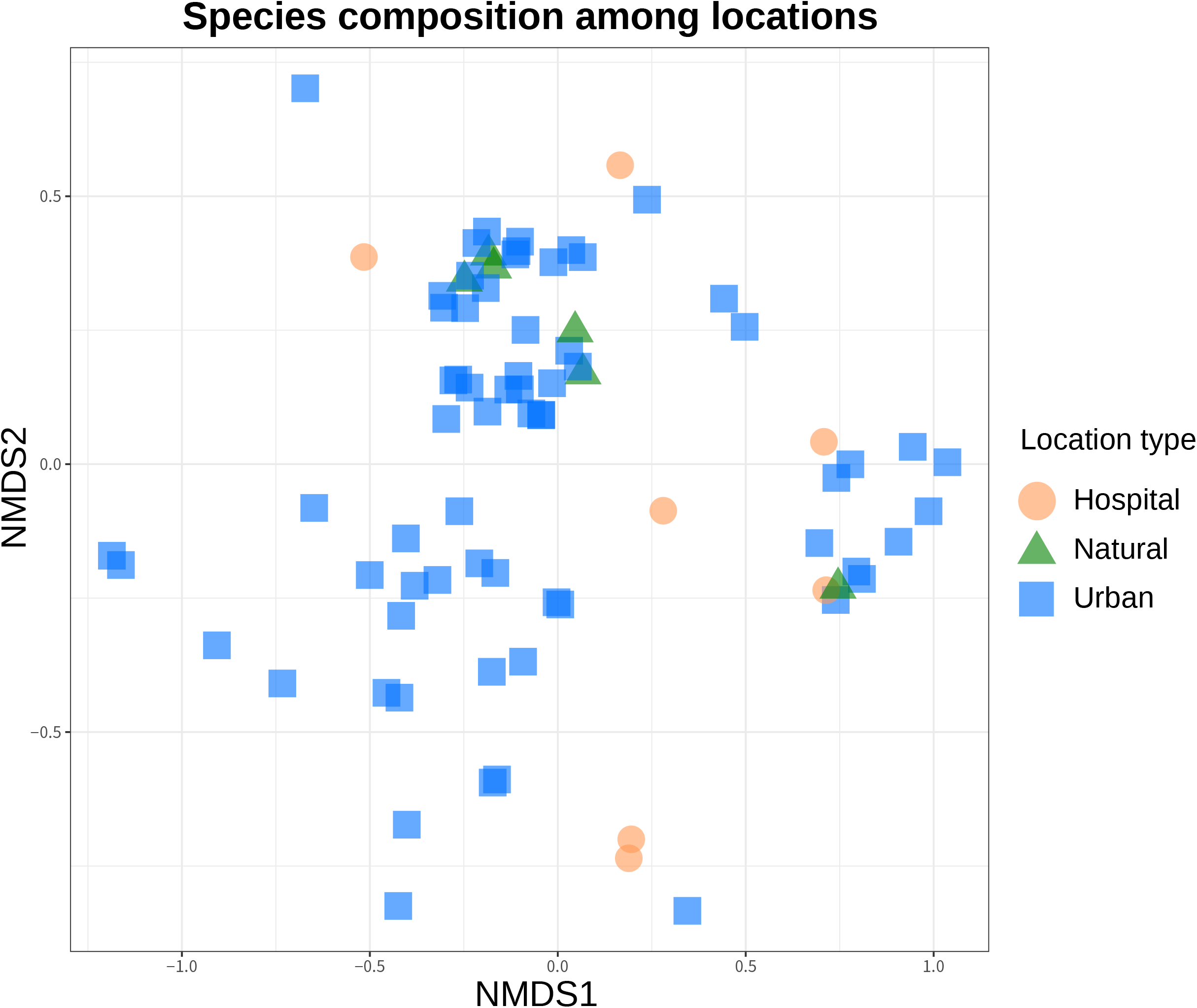
NMDS ordination showing the dissimilarities of species assemblage in soil samples from each location.

General linear model-based analysis indicated an evident effect of location types (i.e. urban, hospital, and natural samples) on the multivariate composition of the recovered *Scedosporium* and *Lomentospora* assemblages (*P=*0.001) (Table 4, model 1). In addition, the univariate test indicated that *S. apiospermum* (*P=*0.012), *S. boydii* (*P=*0.017), and *S. dehoogii* (*P=*0.047) showed moderate to moderate-strong evidence of its preferred distribution in urban regions (including urban and hospital site) (Table 5, model 1). Similar to the result of PERMANOVA analysis, we found no evidence that the recovered fungal composition was explained by location (*P=*0.097) and its interaction term with location type (*P=*0.533) (Table 4, model 2). In the further test using location and sampling unit as factors (model 3), we found moderate-strong evidence of the location effect (*P=*0.007) and strong evidence of the sampling unit effect (*P=*0.001) on the recovered fungal composition. Furthermore, there was strong evidence indicating that *S. apiospermum* is preferentially recovered from certain sampling units (*P=*0.002), and moderate-strong evidence for *S. boydii* (*P=*0.022) and *S. dehoogii* (*P=*0.039)(Table 5, model 3).

**Table 4.**
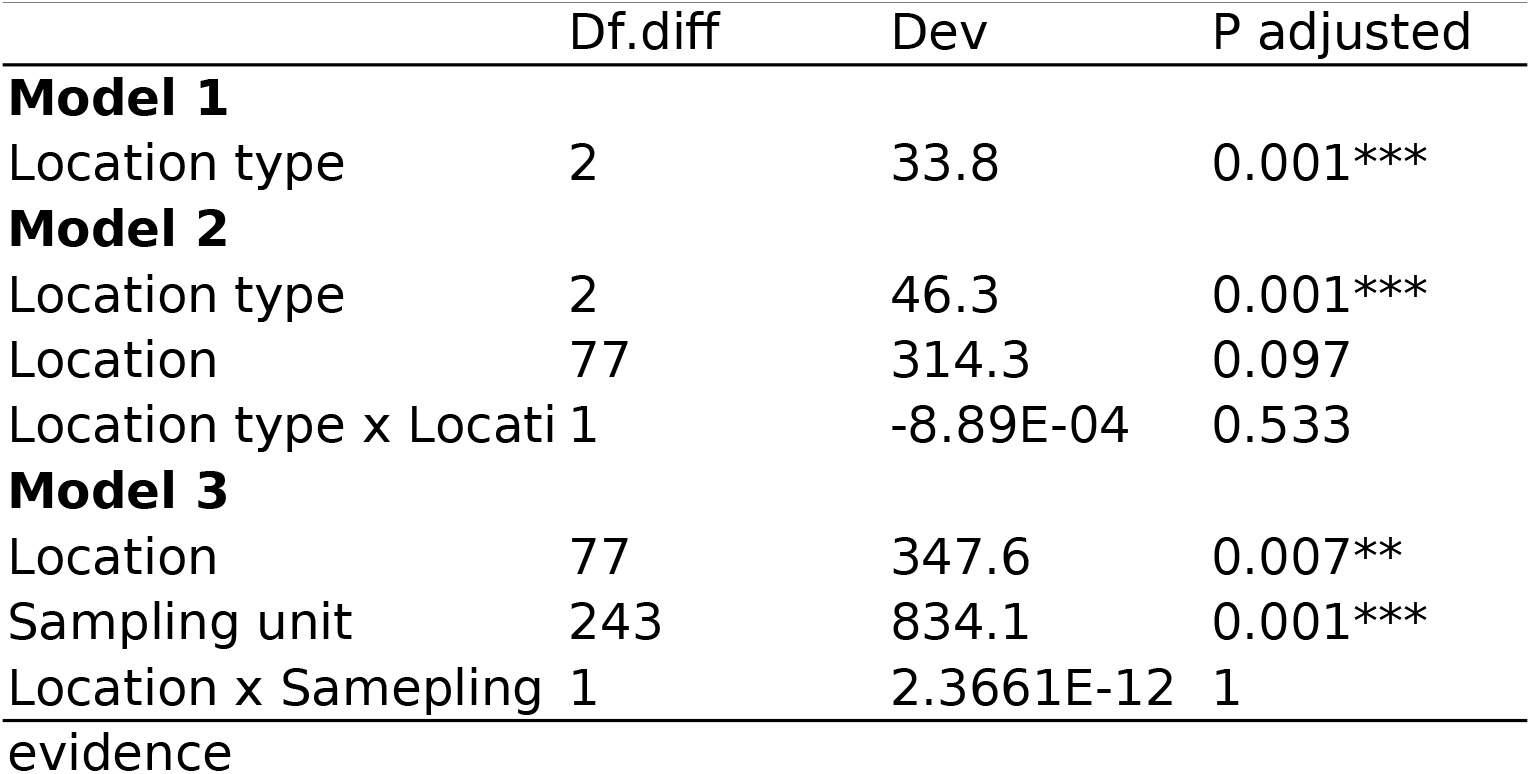
Three models using the general linear-model-based analysis tested the occurrence differences of *Scedosporium* and *Lomentospora* species explained by the effect of location type (model 1); location type, location and the interaction term (model 2); location, sampling unit, and the interaction term (model 3). Degree of freedom, deviance, and alpha-corrected P-values of multivariate tests were shown.

**Table 5.**
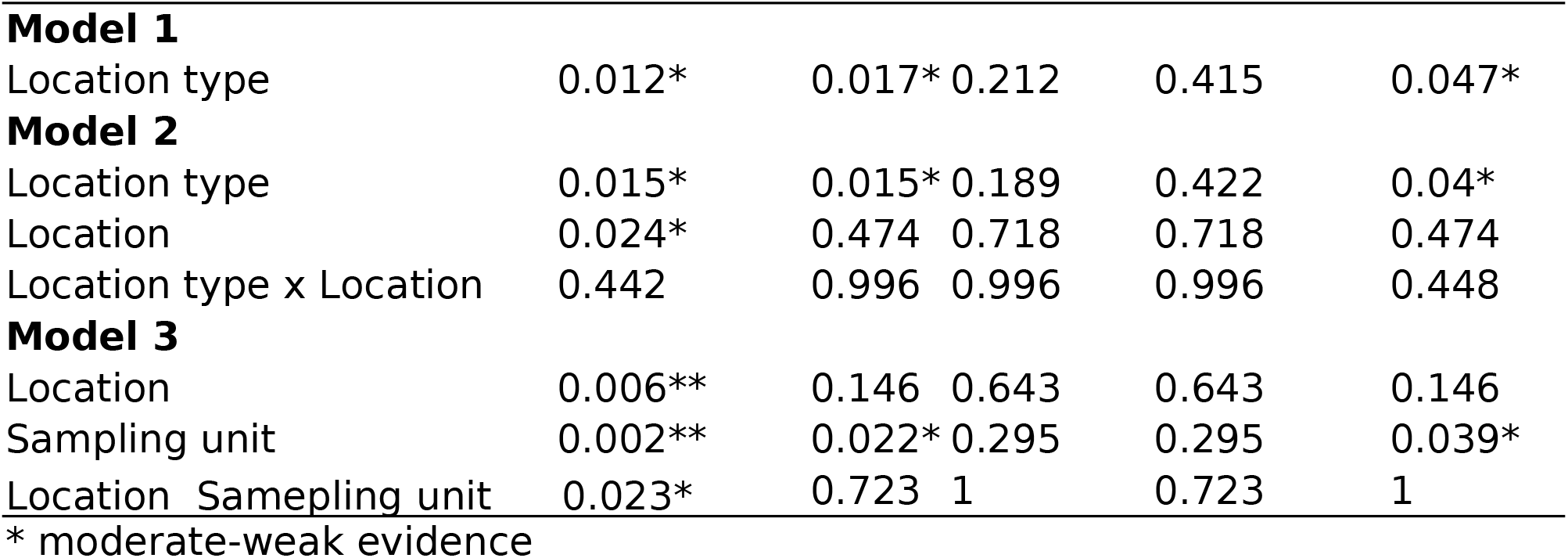
Three models using the general linear-model-based analysis tested the occurrence differences of *Scedosporium* and *Lomentospora* species explained by the effect of location type (model 1); location type, location and the interaction term (model 2); location, sampling unit, and the interaction term (model 3). The alpha-corrected P-values of univariate tests were shown.

## 4. Discussion

*Scedosporium* and *Lomentospora* species were prevalent in soil samples in Taiwan. Their occurrence in the environment were mostly in human disturbed areas, i.e. in urban and hospital samples in the present study. In contrast, natural regions had limited distribution of these fungal species. While detecting *Scedosporium* and *Lomentospora* species in hospital regions in the present study, our results indicated that these hygiene-sensitive areas, in general, possessed no more fungal burden than nearby urban regions; lessening the need of taking a precautionary approach at this point.

### *Scedosporium* and *Lomentospora* are prevalent and are heterogeneously distributed in the environment

Overall, half of the soil samples in the present study yield *Scedosporium* and *Lomentospora* species. Their distribution was not only on the main island, but also on the outlying islands such as Penghu and Kinmen. Although with less density of human population on these islands, we recovered a comparable amount of *Scedosporium* and *Lomentospora* species as that on the main island. Whether the fungal community of these species on the outlying island is endemic or is brought via human activities, and if so the disseminating route, await further investigations.

Distribution of *Scedosporium* and *Lomentospora* species in the environment appear to be highly heterogeneous, even sampling sites in vicinity often yielded varied compositions of these fungal species, as suggested in the homogeneity test (Figure 4). Notably, we did not observe apparent patterns explaining the high fungal burden of these species in certain sampling sites. We found, for example, *S. apiospermum* and *L. prolificans* are abundant in sampling sites in parks where birds are fed and are routinely roosting in; these are places where they may accumulate high nitrogen contents that can encourage the population of *Scedosporium* and *Lomentospora* species (36, 39). Other than this putative factor, we found in some tourist visiting sites (e.g. British Consulate at Takow, Chishang Hakka Cultural Park, and Rushan Visitor Center) and hospitals (e.g. Taichung Veterans Hospital and Taichung Hospital) where they accommodate exceptionally high fungal burden of these species (Figure 3). While again, the distribution of *Scedosporium* and *Lomentospora* species are highly variable in sites of these high fungal burden places. The heterogeneous distribution of these fungal species in the environment invites analysis of the physical and chemical properties of soil samples in future studies.

As in many previous studies, *S. apiospermum* is the most common species in the environment (34, 36, 38–41). The prevalence of this species in the environment has been repeatedly revealed in varied countries across continents and climatic zones, and by using varied selective culturing methods, i.e. dichloran rose bengal chloramphenicol (DRBC) media, *Scedosporium*-selective media (SceSel), and Scedo-Select III culture medium (36, 39, 67). All together, it suggests that *S. apiospermum* appears to be the dominant species in the soil environment globally regardless of geographical and climatic regions. This, in conjunction with their prevalence in human disturbed regions (e.g. sewage, wastewater treatment plants, and landfills), suggests their potential as the indicator species for anthropogenic disturbance or pollution in areas worldwide (23).

### *Scedosporium* and *Lomentospora* have limited distribution in natural regions

In the present study, we isolated *S. apiospermum* from a natural region. As opposed to previous studies, in which no *Scedosporium* and *Lomentospora* species were isolated from the natural/pristine areas (35, 36). The only three cases of recovering *Scedosporium* from natural regions were samples from Argane forests in Souss-Massa, forests in Maamoura, and beaches in Casablanca, all in Morocco (39, 68, 69). These places, however, appeared to exhibit anthropogenic activities, especially in the highly visited Casablanca beach; it may explain the presence of *Scedosporium* species in these regions. Similar to the case in Morocco, the detected sample in the natural region in Taiwan was from Lotus Pond, which is a recreation place with moderate tourist vistings, the low recovery rate and abundance (16 CFU/g) of *Scedosporium* species was therefore plausible. The other five natural places, Cui-Feng, Daxueshan, Baimaoshan, Kuantaoshan, and Liu-Long-Tou, are relatively high-elevated and with lesser tourist visitings; we therefore did not find the fungi in the present study. Intriguingly, few samples from a recreation area (Moon World) with the Badlands landscape exhibit the distribution of *S. apiospermum* (16 CFU/g) and *L. prolificans* (8 CFU/g). Though with low fungal burdens for both species, their presences in such an arid region suggest the tolerance of *Scedosporium* and *Lomemtospora* species to many adverse environments.

### Surveys in different countries revealed varied community profiles of *Scedosporium* and *Lomentospora*

The four clinical relevant species, i.e. *S. apiospermum* (42.5%), *S. boydii* (27.5%), *L. prolificans* (5.6%), and *S. aurantiacum* (8.7%), accounted for 84.3% of the recovered fungal species in the present study; the clinically minor species, *S. dehogii* (15.0%) and *S. haikouense* (0.6%), accounted for the rest recovered isolates. This recovered fungal community profile in Taiwan is different from that in different countries. For example, an environmental survey in Morocco revealed a similar fungal profile of *S. apiospermum* (56%), *S. aurantiacum* (17%), *S. boydii* (18%), and *S. dehogii* (9%) (39); in Thailand revealed a profile of *S. apiospermum* (87%), *S. aurantiacum* (7%), *S. dehogii* (6%) (40); in France revealed a profile of *S. dehogii* (39.4%), *S. aurantiacum* (21.6%), *S. boydii* (19.7%), and *S. apiospermum* (18.9%) (35); in Australia revealed a profile of *S. aurantiacum* (54.6%), *L. prolificans* (43%), *S. boydii* (2.1%), *S. dehogii* (0.3%), and *S. minutispora* (0.004%) (34); and in Austria, a profile of *S. apiospermum* (69.8%), *S. dehoogii* (19.8%), *S. aurantiacum* (6.3%), *S. boydii* (2.2%), and *S. minutispora* (1.9%) was revealed (36).

Intriguingly, the disclosed fungal profile of *Scedosporium* and *Lomentospora* in countries appear to echo the clinical cases attributed to certain species. For example, patients with chronic lung disease in Australia were mostly colonized by *S. aurantiacum* (34). Clinical cases in Austria were mostly attributed to *S. apiospermum* (36). Infections in Taiwan were diagnosed with *S. apiospermum* and *S. boydii* as the culprits (Sun et al. unpublished). While discrepancy happens such as the case in France where *S. boydii* is the major cause of infections but the dominant species was *S. dehoogii* in the environmental study. The exact link between environmental and clinical strains warrants future studies using a more rigorous experimental design coupled with approaches such as molecular population genetics.

---

**Figure S1.**
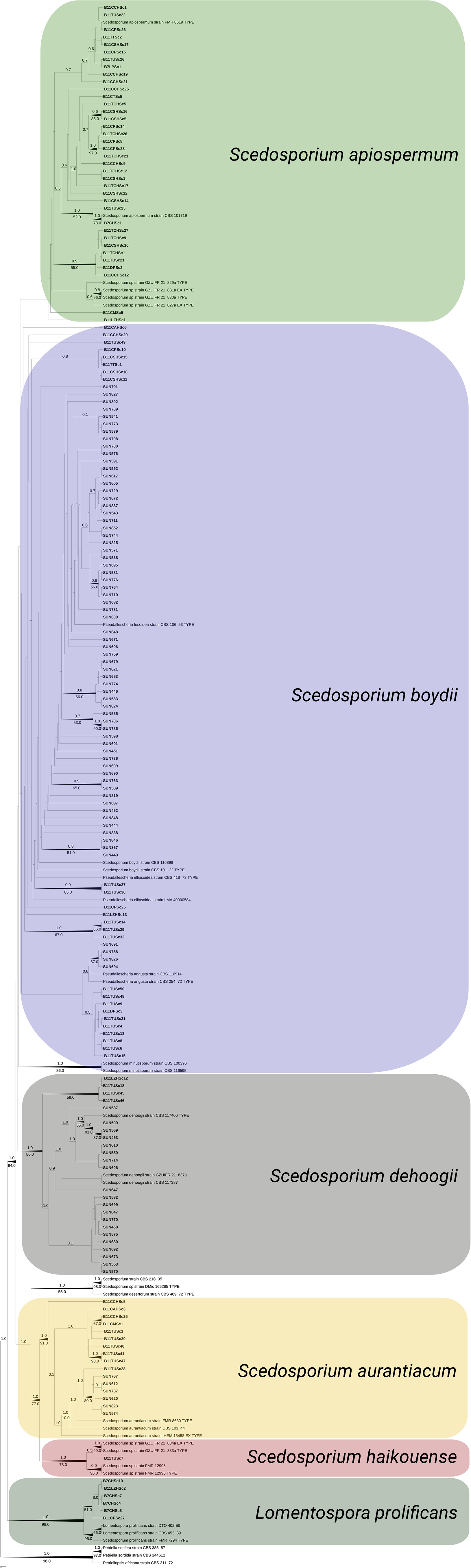
Phylogeny obtained by Maximum Likelihood analysis of the internal transcribed spacer (ITS) sequence dataset showing the phylogenetic relationship of *Scedosporium* and *Lomentospora* species. *Petriellopsis africana* (CBS311.72), *Petriella setifera* (CBS 385.87), and *Petriella sordida* (CBS 144612) were selected as the outgroup. Numbers above nodes are Bayesian posterior probability values; numbers below nodes are maximum likelihood bootstrap values. Type and ex-type species are indicated on the taxon name. Nodes in bold are species isolated in the present study.

**Table S1.**
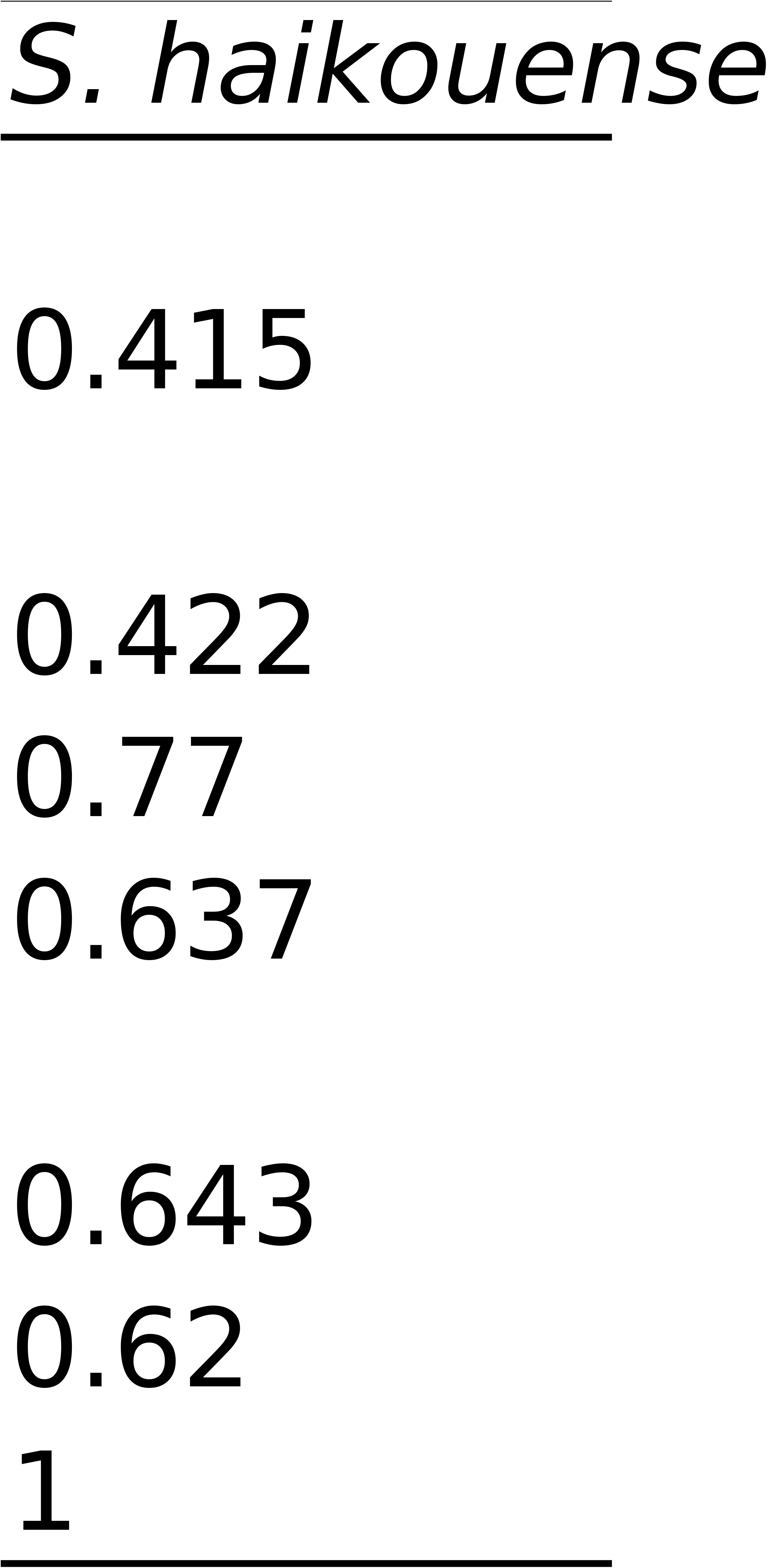
Primer names and their sequences used in the present study.

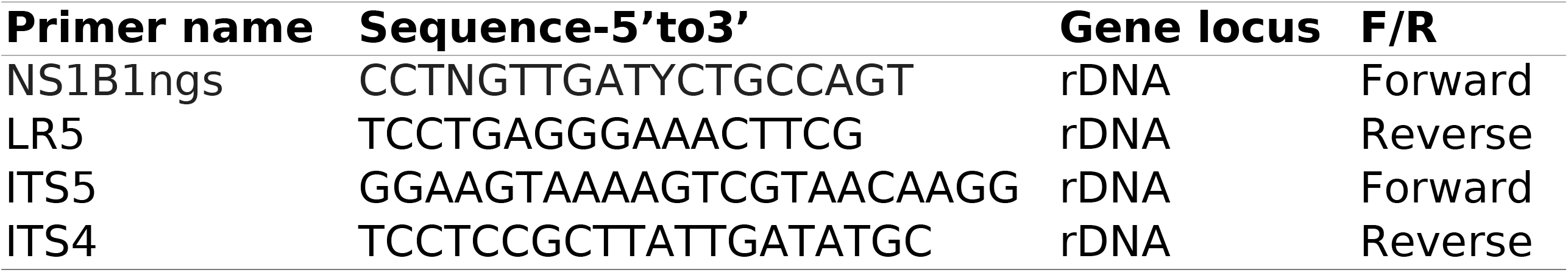

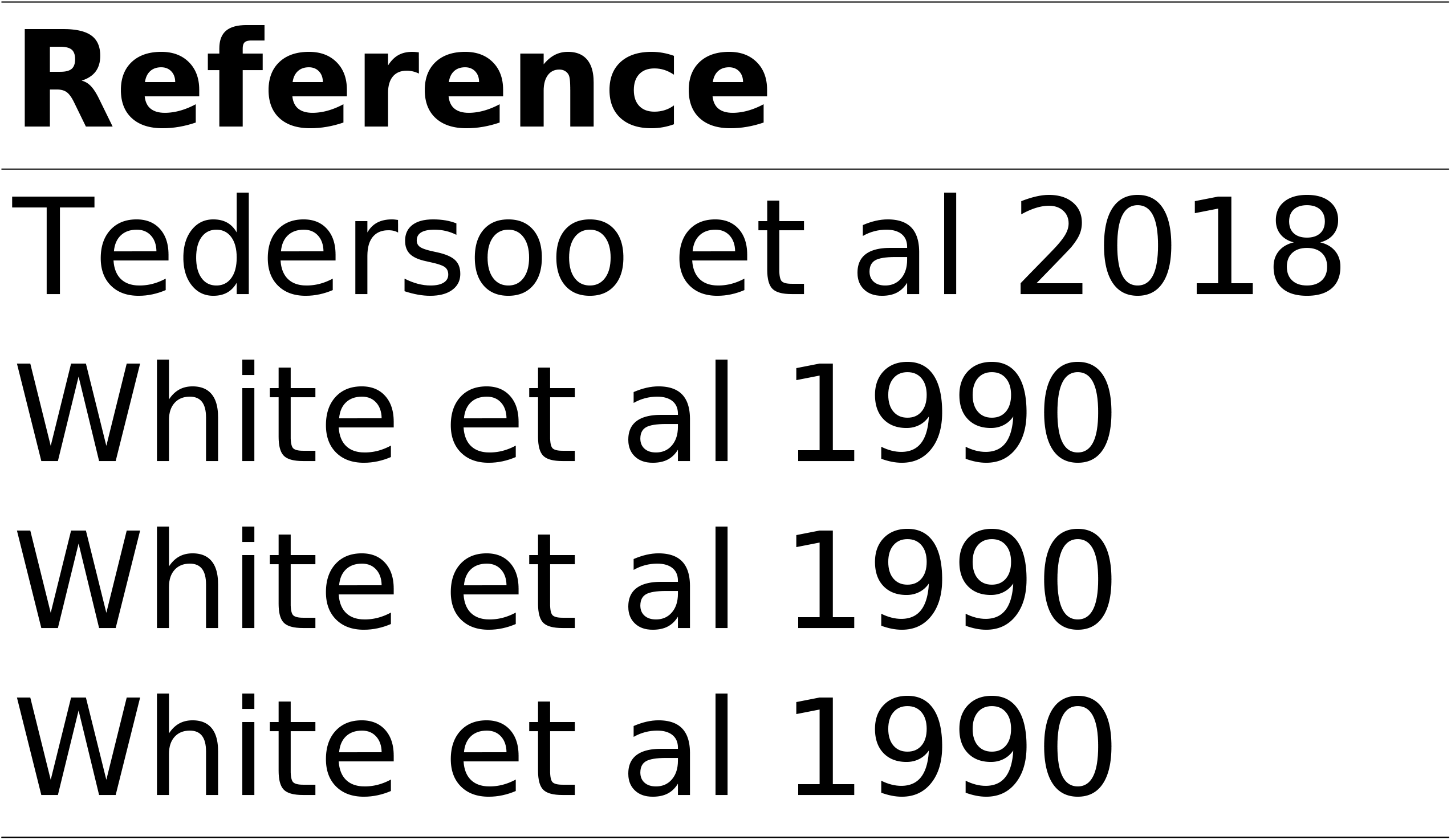

